# Phosphorus is a critical factor of the *in vitro* monoxenic culture method for a wide range of arbuscular mycorrhizal fungi culture collections

**DOI:** 10.1101/2021.09.07.459222

**Authors:** Takumi Sato, Kenta Suzuki, Erika Usui, Yasunori Ichihashi

## Abstract

Establishing an effective way to propagate a wide range of arbuscular mycorrhizal (AM) fungi species is desirable for mycorrhizal research and agricultural applications. Although the success of mycorrhizal formation is required for spore production of AM fungi, the critical factors for its construction in the *in vitro* monoxenic culture protocol remain to be identified. In this study, we evaluated the growth of hairy roots from carrot, flax, and chicory, and investigated the effects of the phosphorus (P) concentration in the mother plate, as well as the levels of P, sucrose, and macronutrients in a cocultivation plate with a hairy root, amount of medium of the cocultivation plate, and location of spore inoculation, by utilizing the Bayesian information criterion model selection with greater than 800 units of data. We found that the flax hairy root was suitable for *in vitro* monoxenic culture, and that the concentration of P in the cocultivation plate was a critical factor for mycorrhizal formation. We showed that an extremely low concentration of P (3 μM) significantly improved mycorrhizal formation for AM fungi belonging to the Glomerales order, while a high concentration of P (30 μM) was suitable for Diversisporales fungi. Therefore, we anticipate that the refining the P concentration will contribute to future culture collections of a wide range of AM fungi.

## Introduction

Arbuscular mycorrhizal (AM) fungi are obligate biotrophs that develop symbiotic relationships with the vast majority of land plants, to create arguably the world’s most prevalent mutualism (Brundrett, 2009; Smith and Read, 2008). Through symbiosis, AM fungi allow plants to obtain soil resources effectively, while receiving carbon in the form of sugars and lipids from the host plants (Luginbuehl et al., 2017). These fungi also confer tolerance against environmental stresses such as water, salinity, and heavy metal levels to the hosts (Augé et al., 2014; Díaz et al., 1996; Porcel et al., 2012), and enhance disease resistance and photosynthetic activity (Boldt et al., 2011; Jung et al., 2012; Pozo & Azcón-Aguilar, 2007). Because of these beneficial effects to plants, substantial efforts have been made to establish ways to propagate AM fungi for agricultural applications (Berruti et al., 2016; Sawers et al., 2008).

AM fungi belong to the subphylum Glomeromycotina and originated in the Ordovician period, approximately 480 million years ago (Redecker et al., 2000; Spatafora et al., 2016). Approximately 342 AM fungal species have been reported and classified into four orders (Diversisporales, Glomerales, Archaeosporales, and Paraglomerales), 11 families, and 43 genera (Redecker et al. 2013, http://www.amf-phylogeny.com/amphylo_species.html). Since the majority are unculturable under *in vitro* conditions, their symbiotic functions in host plants are largely unknown. Since AM fungi and their host plants interact in complex underground networks involving multiple partners, an *in vitro* pure culture method for an isolated AM fungus is required to directly assess the symbiotic interaction between AM fungi and their hosts.

One of the most successful methods of *in vitro* pure culture for AM fungi is the use of Ri T-DNA transformed roots (hairy roots), known as *in vitro* monoxenic culturing (Bécard & Fortin, 1988a; Chabot et al., 1992; Cranenbrouck et al., 2005; Mugnier & Mosse, 1987). In the past three decades, *in vitro* monoxenic culturing with hairy roots has become an important technique for investigating the physiology of AM fungi and their associations with host plants (Fortin et al., 2002). Using this method, spores of the AM fungus *Rhizophagus irregularis* can be propagated *in vitro* for laboratory and greenhouse use and as a commercial inoculum for increasing crop yields (Berruti et al., 2016), such as that of potato (Hijri, 2016), wheat (Al-Karaki et al., 2004), and cassava (Ceballos et al., 2013). Although there have been intensive efforts to improve spore production of AM fungi in the protocols for *in vitro* monoxenic culture (Douds, 2002; Rosikiewicz et al., 2017), the success rate of mycorrhizal formation, which is a prerequisite for spore production, remains to be determined. In particular, there are no known reports assessing the critical factors for mycorrhizal formation in the *in vitro* monoxenic culture protocol for several AM fungal species.

In this study, we investigated six factors (the concentration of P in the mother plate; the concentration of P, sucrose, and macronutrient medium in a cocultivation plate with hairy root; the amount of medium in the cocultivation plate; and the location of spore inoculation) modified by the standard method provided by the Glomeromycota *in vitro* collection (GINCO, http://www.mycorrhiza.be/ginco-bel/) to identify critical factor(s) for mycorrhizal formation in the *in vitro* monoxenic culture. We found that the concentration of P in the cocultivation plate is a critical factor for mycorrhizal formation by regression analysis and showed that extremely low concentrations using flax hairy roots significantly improved the mycorrhizal formation of *R. irregularis*. In addition, we demonstrated that the modification the success rate of mycorrhizal formation for other AM fungi belonging to Glomerales can be higher than that of the standard method. Thus, altering the P concentration of the *in vitro* monoxenic culture method can contribute to a wide range of AM fungi culture collections.

## Materials and methods

### Biological materials

*Agrobacterium rhizogenes*-transformed roots of carrot (*Daucus carota* strain carrot DC2) flax (*Linum* sp. L. strain flax NM) and chicory (*Cichorium intybus* L. strain ChicoryA4NH), known as hairy roots, were obtained from GINCO. All were cultured on a modified Strullu–Romand (MSR) medium used as a cocultivation plate for AM fungi (Cranenbrouck et al., 2005) in Petri dishes (Asnol sterilization Petri dish, GD90-15, 90 mm diameter). The plates for culturing hairy roots were incubated in an inverted position in the dark at 25 °C and maintained by sub-culturing every 4 weeks.

Sterile spore suspensions of the AM fungus *Rhizophagus irregularis* DAOM197198 and the inoculum of National Agriculture and Food Research Organization (NARO) AM fungi *Claroideoglomus etunicatum* MAFF520053 *Scutellospora cerradensis* MAFF520056, *Ambispora callosa* MAFF520057, *Acaulospora longula* MAFF520060, *Gigaspora rosea* MAFF520062, *Acaulospora morrowiae* MAFF520081, *Rhizophagus clarus* MAFF520086, *Paraglomus occultum* MAFF520091, and *Claroideoglomus claroideum* MAFF520092 were obtained from Premier Tech (Quebec, Canada), and NARO Genebank (https://www.gene.affrc.go.jp/index_j.php), respectively. The strains of NARO AM fungi were propagated with Welsh onion (*Allium fistulosum* L. ‘Motokura’), sorghum (*Sorghum bicolor* (L.) Moench. ‘Ryokuhiyou sorugo’), and white clover (*Trifolium incarnatum* L. ‘Dixie’) grown in pots with quartz sand (Mikawakeisa Nomal No.5, Mikawakeisa Corp., Aichi, Japan). Centrifuge tubes (50 mL, Labcon North America, Petaluma, CA, USA) with an 8-mm-diameter hole with cotton (Φ8×25 mm, Safe Basic cotton roll, A.R. Medicom Inc. (Asia) Ltd., Hyogo, Japan) at the bottom were used as pots. Each spore was isolated from each inoculum of AM fungi by wet sieving (500, 100, and 45 μm meshes) and decanting (Gerdemann, 1955). Twenty spores of each AM fungus were placed between 30 g and 35 g of quartz sand in the pot, and 3–4 seeds of each host were sown on the upper layer of the sand. Each pot was irrigated with 10 mL of reverse osmosis water, covered with plastic wrap until seeds were germinated and thinned to three plants per pot. Every other day for 12 weeks after sowing, each pot was irrigated with 10 mL of half-strength Hoagland’s solution containing 50 μM phosphorus (P). At 12 weeks after sowing, irrigation was stopped for the host plants to senesce. Air-dried soil containing each spore of AM fungi was stored at 4 °C before use.

The half-strength Hoagland’s solution contained 126.4 mg KNO3; 295.2 mg Ca(NO_3_)_2_・4H_2_O; 123.3 mg MgSO_4_・7H_2_O; 34 mg KH_2_PO_4_ (corresponding to 50 μM P); 196 mg K_2_SO_4_; 8.2 mg Fe (III)-EDTA; 0.72 mg H_3_BO_3_; 0.45 mg MnSO4・4H_2_O; 0.06 mg ZnSO_4_・7H_2_O; 0.02 mg CuSO_4_・5H_2_O; and 0.0063 mg Na_2_MoO_4_・2H_2_O in 1 L distilled water.

The MSR medium contained 739 mg MgSO_4_・7H_2_O; 76 mg KNO_3_; 65 mg KCl; 4.1 mg KH_2_PO_4_ (corresponding to 30 μM P); 359 mg Ca(NO_3_)_2_・4H_2_O; 0.9 mg calcium panthotenate; 1×10^−3^ mg biotine; 1 mg nicotinic acid; 0.9 mg pyridoxine; 1 mg thiamine; 0.4 mg cyanocobalamine; 1.6 mg Na Fe (III)-EDTA; 2.45 mg MnSO_4_・4H_2_O; 0.29 mg ZnSO_4_・7H_2_O; 1.86 mg H_3_BO_3_; 0.24 mg CuSO_4_・5H_2_O; 0.0024 mg Na_2_MoO_4_・2H_2_O; 0.035 mg (NH_4_)_6_Mo_7_O_24_・4H_2_O; and 10,000 mg sucrose, solidified with 4,000 mg Phytagel in 1 L distilled water.

All chemicals were provided by Wako Pure Chemical Corp (Osaka, Japan), except for FE(III)-EDTA, Na Fe (III)-EDTA, and Phytagel, which were received from Sigma-Aldrich (USA, MA, Burlington).

### Experiment 1—the evaluation of hairy root growth

Experiment 1 was designed to compare the growth of hairy roots of carrot, flax, and chicory. Root fragments of 1 cm length were cut from the tips of hairy roots after 3 weeks of subculture, transferred onto the MSR medium, and incubated in the dark at 25 °C for 4 weeks. The flesh weights of the hairy roots were then measured (n = 12).

### Experiment 2—the evaluation of the success rate of AM fungal colonization in different culture conditions

Experiment 2 was designed to evaluate different culture conditions and their success rate for AM fungal colonization using flax hairy roots and *R. irregularis* DAOM197198. A list of all culture conditions evaluated in Experiment 2 is presented in Table 1. Using the same conditions as Experiment 1 for a control, 1 cm-long flax hairy root fragments were transplanted onto the MSR medium and inoculated with approximately 100 spores of *R. irregularis* DAOM197198 at a distance of 2.5 cm from the root fragment, and incubated in the dark at 25 °C. The standard culture conditions were modified in the following order: mother plate P concentration (3 and 30 μM), macronutrients in the MSR medium (the original and half of the concentration of N, P, K, Mg, Ca, and S in the MSR medium), P concentration (0.3, 3, 15, and 30 μM) in the MSR medium, sucrose concentration (10 and 20 g/L) in the MSR medium, amount of MSR medium (10, 20, and 40 mL/plate), and location of the spore inoculation (distances of 1, 1.5, 2, 2.5, and 3 cm away from the root tip). Five plates were prepared as replicates, and 2–15 replicates for each condition were tested (Table 1). The success rate of AM fungal colonization was determined by the presence or absence of elongation of the extraradical hyphae (EH) and spore formation (SF) at 30 and 60 days after inoculation. In addition, the number of spores in 3 and 30 μM P in the MSR medium and in the 20 and 40 mL samples of the MSR medium were counted at 11 months after inoculation.

**Table 1.** List of all culture conditions evaluated in this study. Bold indicates culture conditions of GINCOs standard protocol.

| Experimenter | Mother P | Essential elements | P concentration | Sucrose concentration | Amount of MSR medium | Location of spore inoculation | Reprication |
| --- | --- | --- | --- | --- | --- | --- | --- |
| | $\mu\text{M}$ | fold | $\mu\text{M}$ | $\text{g L}^{-1}$ | $\text{mL plate}^{-1}$ | cm | |
| A | **30** | **1** | **30** | **10** | **20** | **1** | **10** |
| A | 30 | 1 | 30 | 10 | 20 | 1.5 | 10 |
| A | 30 | 1 | 30 | 10 | 20 | 2 | 10 |
| A | 30 | 1 | 30 | 10 | 20 | 2.5 | 10 |
| A | 30 | 1 | 30 | 10 | 20 | 3 | 10 |
| A | 30 | 1 | 30 | 10 | 10 | 2.5 | 5 |
| A | 30 | 1 | 30 | 10 | 40 | 2.5 | 5 |
| A | 30 | 1 | 30 | 10 | 20 | 2.5 | 5 |
| A | 30 | 0.5 | 15 | 10 | 20 | 2.5 | 5 |
| A | 30 | 0.5 | 3 | 10 | 20 | 2.5 | 5 |
| A | 3 | 1 | 30 | 10 | 20 | 2.5 | 5 |
| A | 3 | 0.5 | 15 | 10 | 20 | 2.5 | 5 |
| A | 3 | 0.5 | 3 | 10 | 20 | 2.5 | 5 |
| A | 3 | 1 | 30 | 10 | 40 | 2.5 | 2 |
| A | 3 | 0.5 | 15 | 10 | 40 | 2.5 | 2 |
| A | 3 | 0.5 | 3 | 10 | 40 | 2.5 | 2 |
| B | **30** | **1** | **30** | **10** | **20** | **1** | **5** |
| B | 30 | 1 | 30 | 10 | 20 | 2.5 | 15 |
| B | 3 | 1 | 30 | 10 | 20 | 2.5 | 5 |
| B | 30 | 1 | 3 | 10 | 20 | 2.5 | 5 |
| B | 3 | 1 | 3 | 10 | 20 | 2.5 | 5 |
| B | 30 | 1 | 0.3 | 10 | 20 | 2.5 | 5 |
| B | 3 | 0.5 | 3 | 10 | 20 | 2.5 | 5 |
| B | 30 | 0.5 | 3 | 10 | 20 | 2.5 | 5 |
| B | 30 | 1 | 30 | 20 | 20 | 2.5 | 5 |
| B | 30 | 0.5 | 3 | 20 | 20 | 2.5 | 5 |

### Experiment 3—the evaluation of the success rate of colonization for different species of AM fungi in the optimized culture condition

Experiment 3 was designed to evaluate whether the concentration of P in the MSR medium is a critical factor based on Experiment 2 using *R. irregularis*, for the success rate of colonization of different AM fungal species. Spores of the NARO AM fungal species were collected from the propagated inoculum by wet sieving and the decanting technique described above. The collected spores were surface-sterilized using a modified method described by Bécard and Fortin (1988). Briefly, spores were sonicated with sterile 0.05% Tween 20 solution three times, soaked in 2% (w/v) chloramine T solution for 10 min, and rinsed three times with sterile 0.05% (w/v) Tween 20 solution. The surface-sterilized spores were stored in a sterile solution containing 0.02% (w/v) streptomycin and 0.01% (w/v) gentamycin at 4 °C until use. Each spore was inoculated onto the MSR medium with 1-cm-long flax hairy root fragments, inoculated with 5 spores at a distance of 2.5 cm from the root fragment, and incubated in the dark at 25 °C. The germinated spores were transferred from MSR medium with a scalpel and tweezers to 3 or 30 μM P MSR medium. As previously mentioned, the success rates of AM fungal colonization and the presence or absence of elongation of EH and SF for different AM fungal species were determined at 30, 60, and 90 days after inoculation.

### Statistical analysis

All statistical analyses were performed using R version 3.4.0. The data on the success rate of AM fungal colonization for all treatments were pooled and used for logistic regression analysis with model selection using the ‘bestglm’ utility of the R package (McLeod et al., 2020) to identify the main factors that contribute to the success rate of AM fungal colonization, and the mathematical functions that describe the methods these factors use to explain the success rate. By default, ‘bestglm’ uses the Bayesian information criterion (BIC) for model selection. Since BIC is appropriate for controlled experiments with a limited number of important explanatory variables (Aho et al., 2014), it was deemed acceptable for identifying relevant common factors among the different criteria for the success of AM fungal colonization. To examine the differences among experimental groups, data were analyzed with Welch’s *t-*test, Tukey’s honest significant difference test, and the Games-Howell test (*P* < 0.05).

## Results

### Flax hairy root is suitable for *in vitro* monoxenic culture of AM fungi

Carrot hairy roots have been widely used for *in vitro* monoxenic cultures of AM fungi (Cranenbrouck et al., 2005; Fortin et al., 2002; Kokkoris & Hart, 2019a). Recently, GINCO has provided flax and chicory hairy roots in addition to carrot hairy roots, and we assessed the flesh weight for the three varieties. The results showed that the flax hairy roots had significantly higher growth rates than chicory and carrot hairy roots during the four weeks of incubation (9.44- and 1.88-times, respectively) (Fig 1A). In addition, the flax hairy root revealed the lowest coefficient of variation in dry weight (Fig 1B). This result indicates that flax hairy roots are suitable for *in vitro* monoxenic cultures of AM fungi because of their vigorous growth and robustness against experimental conditions.

**Figure 1.**
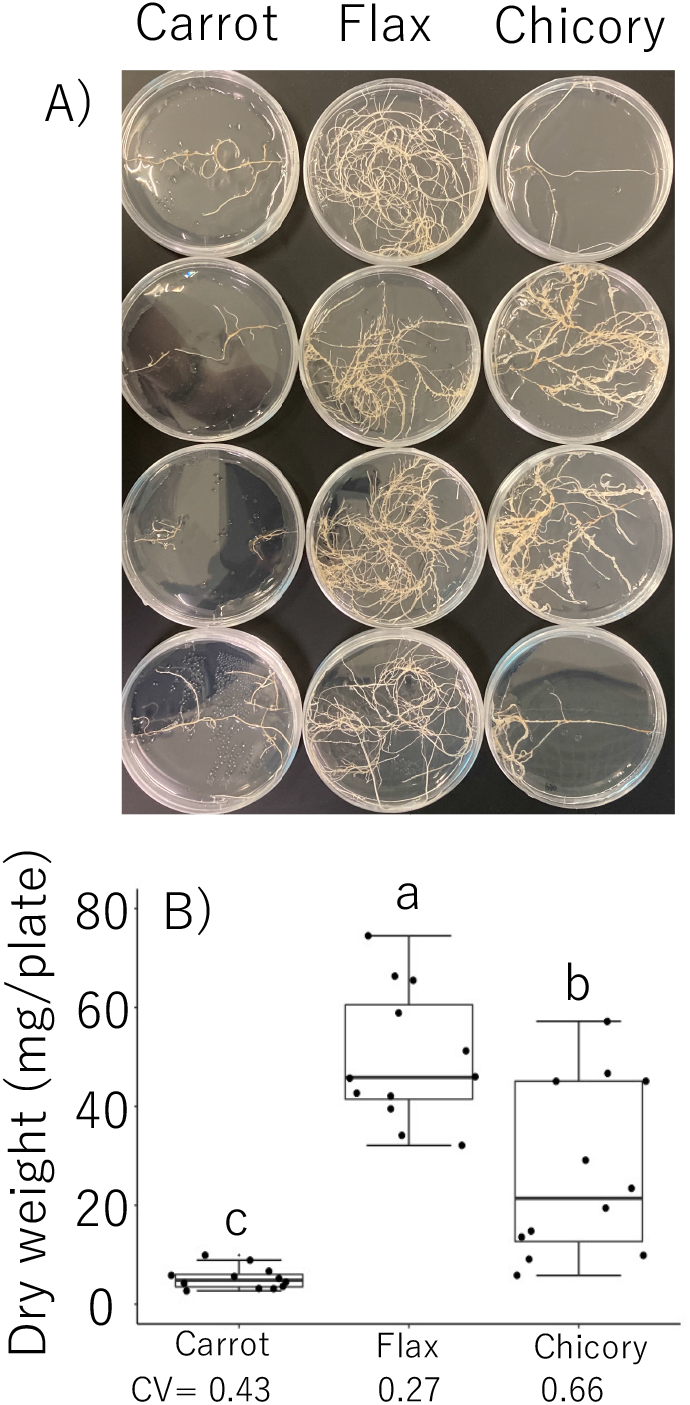
Evaluation of growth of hairy roots derived from different plant species. Gross morphology (A) and dry weight (B) from hairy root of carrot (*Daucus carota* strain carrotDC2), flax (*Linum* sp. strain flax NM), and chicory (*Cichorium intybus* strain ChicoryA4NH) at 4 weeks after incubation are shown. Error bars and CV indicate the standard error and coefficient of variance, respectively (n=12). Different letters indicate significant differences as determined by Tukey’s honestly significant difference test (*P* <0.05).

### Success rate of AM fungal colonization in different culture conditions

To identify the critical factor(s) for mycorrhizal formation in the *in vitro* monoxenic culture using flax hairy roots, we investigated the effects of different culture conditions (the concentration of P in the mother plate, the concentration of P, sucrose, and macronutrients in the cocultivation MSR medium with hairy root, the amount of cocultivation MSR medium, and the location of spore inoculation) on the success rate of AM fungal colonization, through the presence or absence of elongation of extraradical hyphae (EH), and the formation of spores (SF). We then compared the results to the standard method provided by GINCO (Fig. 2), and significant differences (Welch’s *t-*test for two parameters; Games-Howell test for more than three parameters, *P* < 0.05) were detected for all parameters except for sucrose concentration at 30 and 60 DAI and location of spore inoculation at 60 DAI (Fig. 2). To estimate the significant parameters for predicting the success rate of AM fungal colonization, we performed a regression analysis using all the data collected in this study. For each of the binary variables based on the four success criteria as response variables, we identified a combination of explanatory parameters that minimized the BIC and their relative contributions (Table 2). The only factor included in all models was the concentration of P, and it had the largest contribution in all cases. In addition, the best model was obtained when the concentration of P was squared after non-linear logarithmic transformation in all cases (Table 3). Given that the coefficient of log(x)^2^ (x represents the concentration of P) was negative for all models, the colonization success rate peaked at a low P concentration (Fig. 3). The modification with 3 μM P in the MSR medium significantly improved the success rate of AM fungal colonization by 2.05–3.25 times for EH and 1.50–8.00 times for SF compared with the standard method, suggesting that 3 μM P in the MSR medium was an optimal concentration for the success rate of *R. irregularis* colonization (Fig. 2).

**Figure 2.**
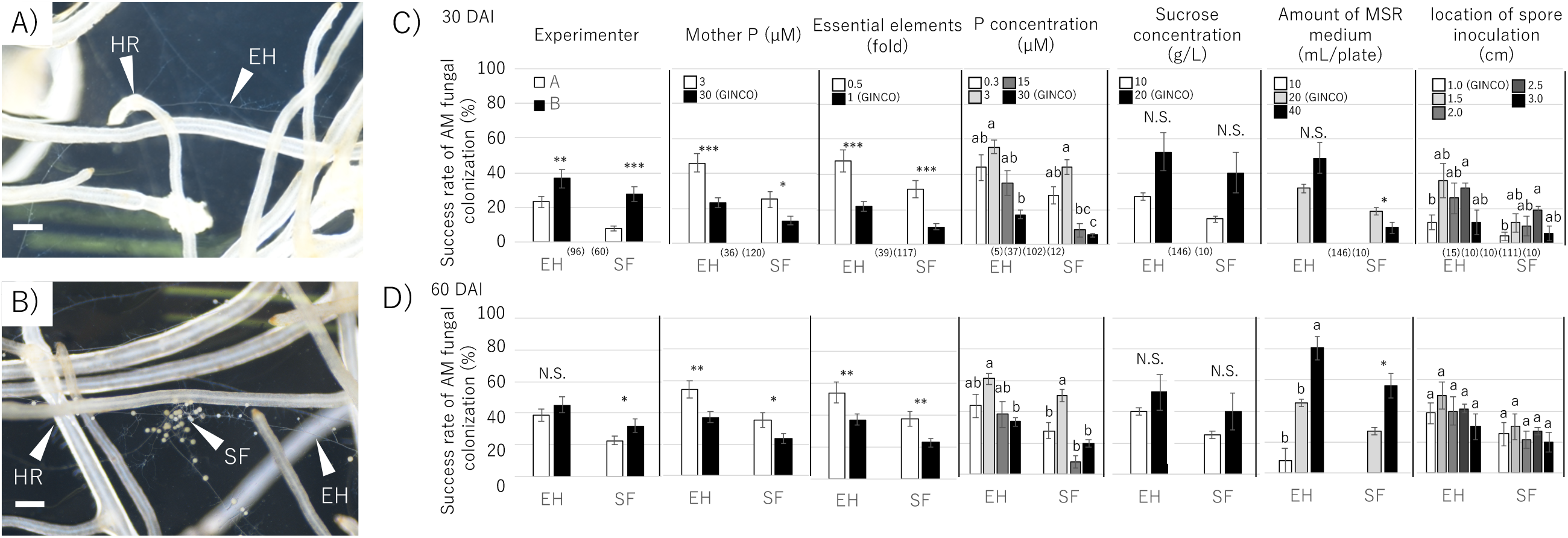
Evaluation of the success rate of AM fungal colonization in different culture conditions. Elongation of extraradical hyphae (EH) from hairy roots (HR) of Flax (*Linum* sp. MUCLflax NM, A), and spore formation (SF, B). White bars indicate 500 μm. Success rate of arbuscular mycorrhizal (AM) fungal colonization, calculated by elongation of extraradical hyphae (EH) and spore formation (SF) of each plate, for each treatment at 30 (C) and 60 (D) days after inoculation (DAI). Error bars indicate standard errors. Different letters indicate significant differences as determined by the Games-Howell test (*P* <0.05). Asterisks indicate significant differences by Welch’s *t-*test (***, *P* < 0.001; **, *P* < 0.01; *, *P* < 0.05).

**Table 2.** BIC of the best model and the contribution of selected factors. The contribution of an explanatory variable is shown as the amount of decrease in BIC when that variable is excluded. A blank space means that the factor was not selected as the explanatory variable.

| | | BIC | △BIC | | | | | |
| --- | --- | --- | --- | --- | --- | --- | --- | --- |
| Days after inoculation |  |  | Mother P | Macronutrient medium | Medium P | Medium C | Amount of medium | Inoculated position |
| 30 | EH | 819.9 | 10.7 |  | 60.5 | 10 |  |  |
|  | SF | 538.6 |  |  | 121.9 | 8.9 |  |  |
| 60 | EH | 995.8 |  |  | 41.7 |  | 21.6 | 6.6 |
|  | SF | 824.2 |  |  | 56.4 |  | 24.7 | 7.8 |

**Table 3.** Relationship between mathematical transformation of P concentration and BIC, calculated by elongation of extraradical hyphae (EH) and spore formation (SF) of each plate, for 0.3, 3, 15, and 30 μM P treatments at 30 and 60 days after inoculation (DAI). The numbers represent BIC. In the first column, x represents no processing, x^2^ represents squaring, log(x) represents logarithmic transformation, and log(x)^2^ represents squaring after logarithmic transformation.

|  | 30 DAI |  | 60 DAI |  |
| --- | --- | --- | --- | --- |
|  | EH | SF | EH | SF |
| x | 836.4 | 547.5 | 1021.5 | 853.2 |
| $x^2$ | 837.0 | 553.9 | 1023.3 | 859.5 |
| $\log(x)$ | 856.1 | 576.7 | 1029.0 | 860.5 |
| $\log(x)^2$ | 835.8 | 541.7 | 1019.4 | 847.1 |

**Figure 3.**
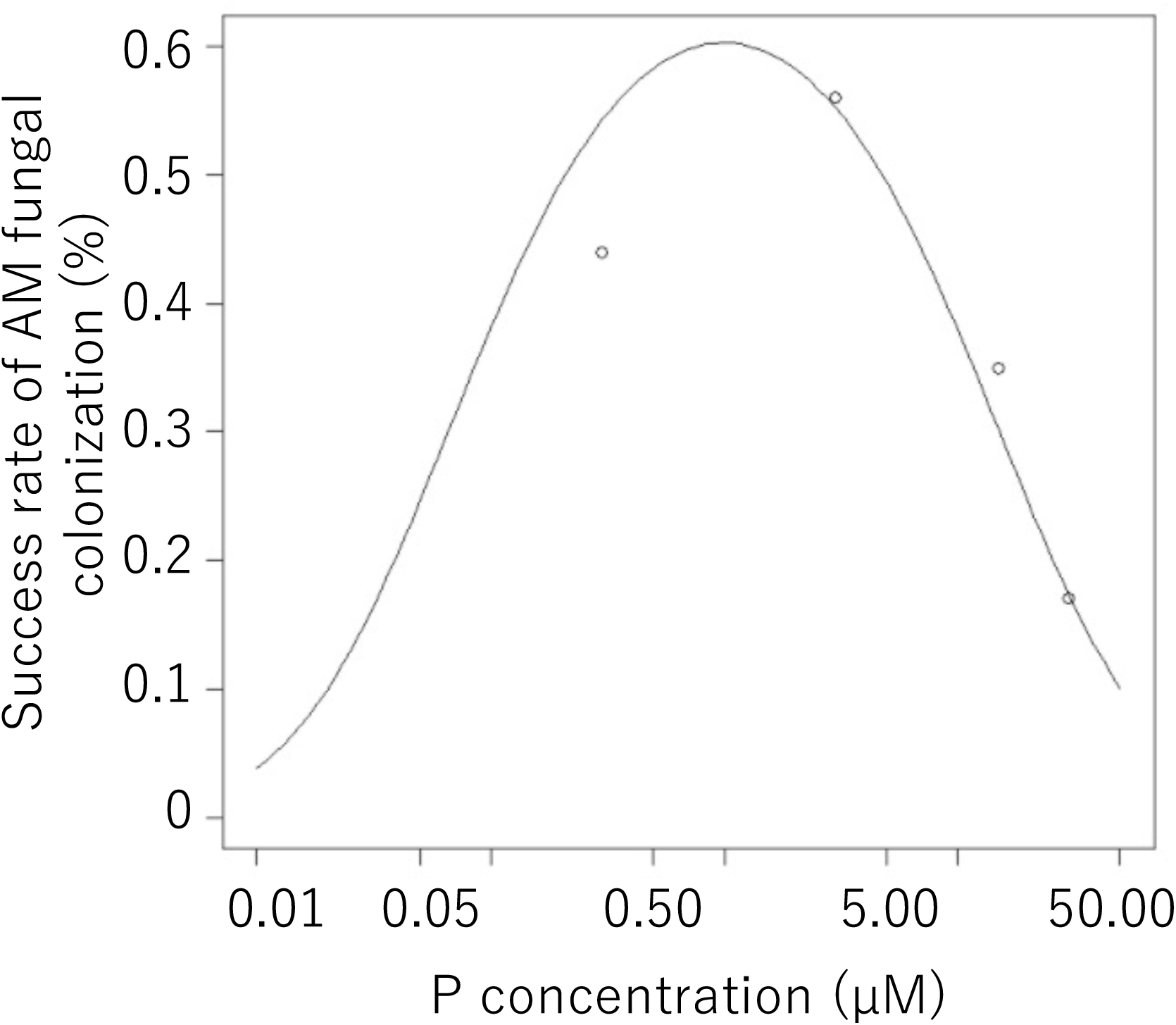
-log (x)^*2*^ model calculated from success rate of arbuscular mycorrhizal (AM) fungal colonization of *Rhizophagus irregularis* DAOM197198, calculated by elongation of extraradical hyphae (EH) for 0.3, 3, 15, and 30 μM P at 30 days after inoculation in Experiment 2.

To validate the improvement of AM fungal colonization prior to sporulation, we observed that the conditions of 3 μM P in the MSR medium produced 2.3 times more spores than that of the control (Welch’s *t-*test, *P* < 0.001, Fig. 4). However, the increased amount of the MSR medium (40 mL), which presented the second greatest contribution in the model selection (Table 2), produced fewer spores than that of the control (20 mL, Fig. 4), which might be because the increased growth of hairy roots provided more opportunities for AM fungal colonization, but the medium did not contain sufficient factors for sporulation.

**Figure 4.**
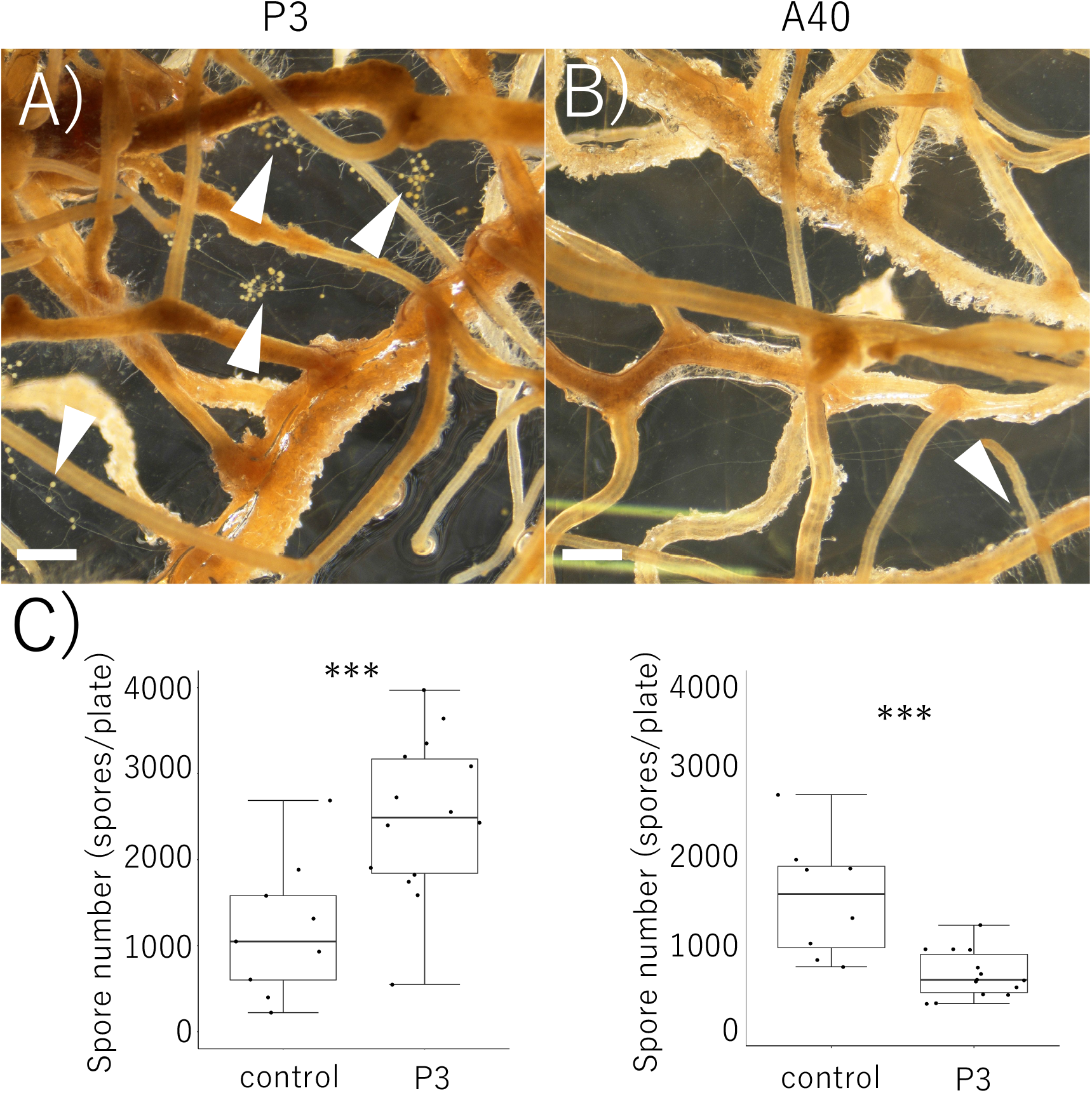
Evaluation of the spore formation in different culture conditions. Spore formation *Rhizophagus irregularis* DAOM197198 P3 (A) and 40 mL (B) treatments in Experiment 2, and number of spores of both treatments (C) at 11 months. Error bars indicate standard errors. White bars indicate 500 μm, and white arrows indicate spores formed in the modified Strullu-Romand medium. Asterisks indicate significant differences by Welch’s *t-*test (***, *P* < 0.001).

### Success rate of colonization for different species of AM fungi in the optimized culture condition

To test whether the modification of P in the MSR medium improved the success rate of AM fungal colonization by other strains, *C. etunicatum* MAFF520053, *S. cerradensis* MAFF520056, *Am. callosa* MAFF520057, *Ac. longula* MAFF520060, *G. rosea* MAFF520062, *Ac. morrowiae* MAFF520081, *R. clarus* MAFF520086, *P. occultum* MAFF520091, *and C. claroideum* MAFF520092 were cultured under the modified conditions (3 μM P) and the standard method (30 μM P). The low concentration of P significantly improved the success rate of AM fungal colonization for *C. etunicatum* MAFF520053 (from 0% to 20% in EH at 60 DAI; Table 4). The same tendency was observed for *Am. callosa* MAFF520057 (from 0% to 12% in EH at 30 DAI; Table 4) and *R. clarus* MAFF520086 (from 0% to 28% in EH and from 0% to 16% in SF at 30 DAI; from 20% to 44% in EH and from 20% to 40% in SF at 60 DAI; and from 24% to 44% in EH and from 24% to 40% in SF at 90 DAI; Table 4). In the case of *G. rosea*, we observed the opposite trend (from 100% to 60% in EH at 30 and 60 DAI; Table 4).

**Table 4.**
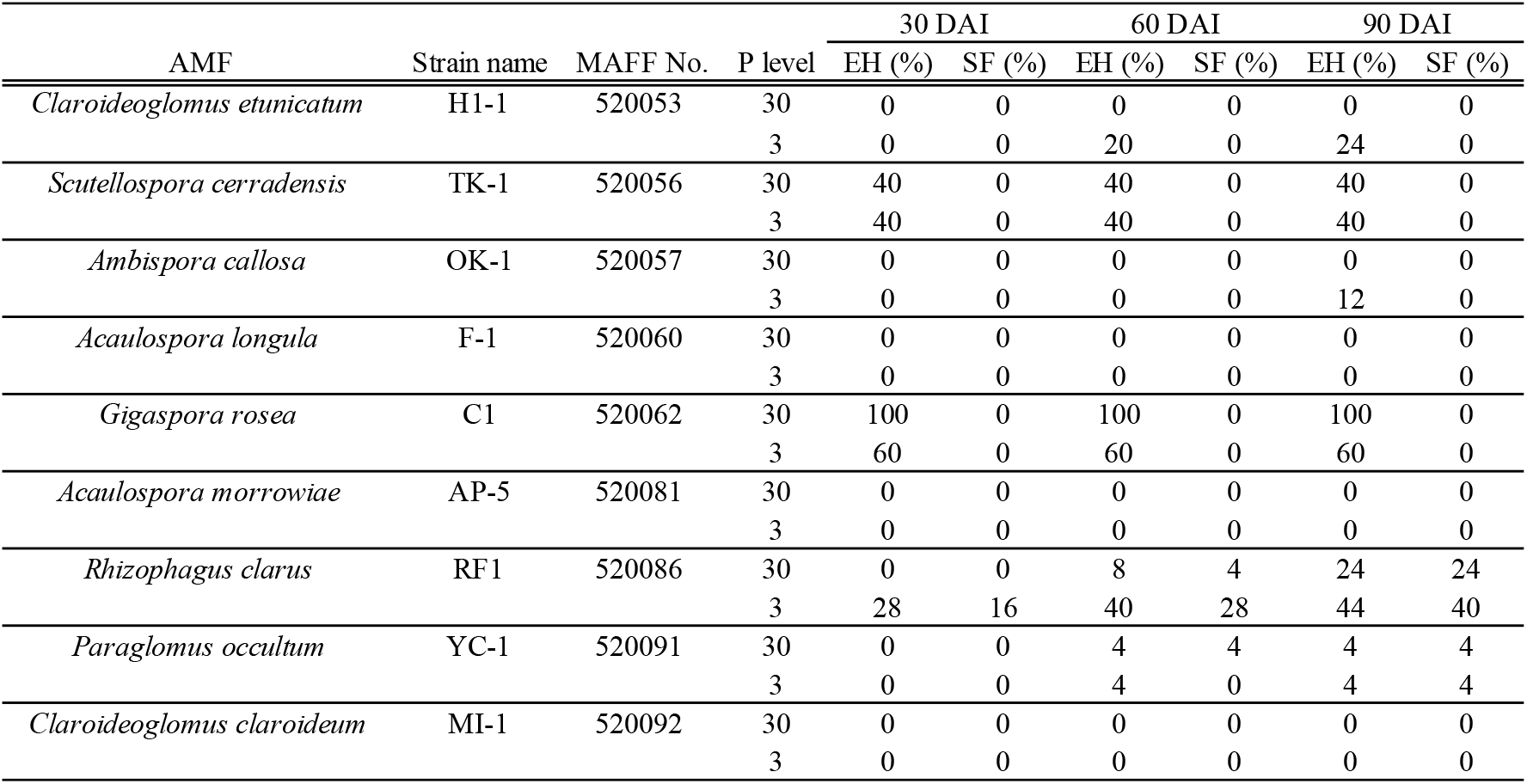
Success rate of arbuscular mycorrhizal (AM) fungal colonization of 9 AMF, calculated by elongation of extraradical hyphae (EH) and spore formation (SF) of each plate, for 30 and 3 μM P treatments at 30, 60, and 90 days after inoculation (DAI).

To validate the improvement of AM fungal colonization leading to sporulation in *R. clarus*, the number of spores of *R. clarus* MAFF520086 was measured only on the spore-forming plates (the number of spore-forming plates in P30 and P3 were 6 and 10, respectively) and found that 3 μM P revealed 3.44 times more spore formation compared to that of 30 μM P (Welch’s *t-*test, *P* < 0.001, Fig. 5).

**Figure 5.**
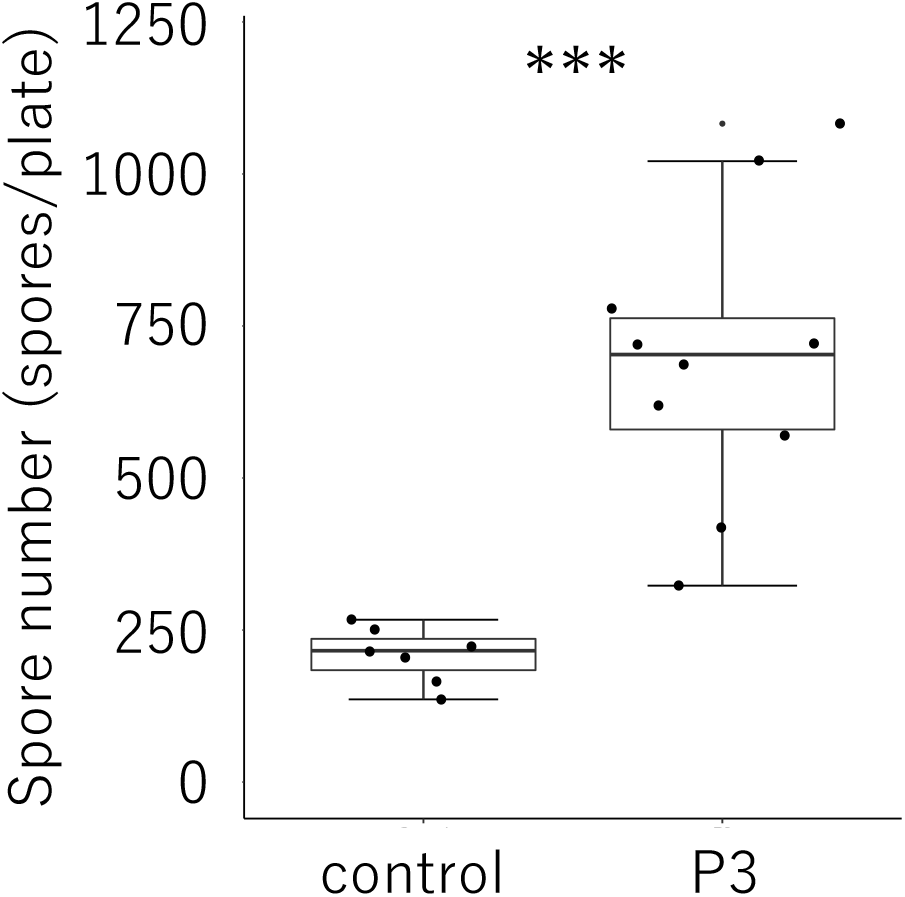
Evaluation of the spore formation in the optimized culture conditions. Number of spores of *Rhizophagus clarus* MAFF 520086 in control (30 μM P) and P3 treatments in Experiment 3 at 90 days after inoculation. Error bars indicate standard errors. Asterisks indicate significant differences by Welchʼs *t*-test (p<0.001 =***).

## Discussion

Over the past three decades, there have been extensive efforts to improve *in vitro* monoxenic culture using carrot hairy roots for AM fungi, with only P-free or 30 μM P as the low-P treatment (Bécard & Fortin, 1988b; Declerck et al., 1998; Douds, 2002; Karandashov et al., 2000; Olsson et al., 2002; Rosikiewicz et al., 2017). Our previous study showed that an extremely low concentration of P (3 μM) in the cocultivation plate using flax hairy roots increased AM fungal colonization (Sato et al., 2019), but the success rate of mycorrhizal formation, which is a prerequisite for spore production, remained to be assessed. This study screened critical factors for the success rate of mycorrhizal formation and found that the P concentration in the cocultivation plate is a critical factor for the mycorrhizal formation of several AM fungi.

Glomerales (i.e., *Rhizophagus irregularis* DAOM197198, *Claroideoglomus etunicatum* MAFF520053, and *Rhizophagus clarus* MAFF520086) showed higher success rates of AM fungal colonization at lower concentrations of P (Table 4). In addition, Archaeosporales (i.e., *Am. callosa* MAFF520057) developed extraradical hyphae only at lower concentrations of P (Table 4). In contrast, Diversisporales (i.e., *Scutellospora cerradensis* MAFF520056 and *Gigaspora rosea* MAFF520062) may prefer higher P concentrations in the MSR medium for the success rate of AM fungal colonization. Although colonization of *Gigaspora margarita*, Diversisporales is enhanced under low phosphorus conditions (Gutjahr et al., 2009; Navazio et al., 2007; Tawaraya et al., 1998) similar to Glomerales, further studies are required. The optimal concentration of P may vary depending on the phylogenetic lineage, since P concentration affects AM fungal colonization and sporulation in a species-specific manner in pot cultures (Silva et al., 2005; Sylvia & Schenck, 1983).

Replacement of the cultivation medium and re-supply of glucose increased the spore production of *Rhizophagus irregularis* (Douds, 2002). Given that the increased sucrose concentration in the cocultivation medium at the beginning of culture did not affect the success rate of mycorrhizal formation in this study (Fig. 2 and Table 2), replenishment of the carbon source after establishment of the plant-AM fungi symbiotic connection would be critical for mycorrhizal formation. Although the transition from the elongation of EH to SF is a gradual continuous change in *R. irregularis*, we observed that the P concentration in the mother plate affected the success rate of EH, but not that of SF, while P and sucrose in the cocultivation medium, which were transferred from the mother plate, are important to the success rate of SF (Table 2). These data suggest that AM fungi should have a mechanism to sense external nutritional conditions, including host status, to regulate their stage transition after colonization.

AM fungi also control the transfer of nutrients depending on plant carbon availability (Hammer et al., 2011), and host plants can distinguish the AM fungi that provide greater P to the host and present them with more carbon sources (Kiers et al., 2011). This bidirectional control for nutrient transfer in plant-AM fungi is described by a theoretical framework reflecting experimental data (Jolicoeur et al., 2002). The development of an effective way to propagate AM fungi must maintain the optimal balance in material transfer to direct the spore production.

The ‘minimal (M)’ medium was developed and subsequently modified for *in vitro* monoxenic culture for AM fungi (Bécard & Fortin, 1988; Pawlowska et al., 1999), but the assessment of different concentrations of P in the medium have been limited. This study showed that a simple modification in the concentration of P in the cocultivation plate greatly improved the success rate of mycorrhizal formation of AM fungi. This modification can be utilized in different types of culture systems (Rosikiewicz et al., 2017), and can enhance AM fungi propagation. Given that long-term *in vitro* propagation has the potential to promote the artificial evolution of AM fungi (Kokkoris & Hart, 2019a, 2019b), the concentration of P would be one of the most important for selective pressure targets for evolution, and an optimal concentration of P similar to the natural habitat would be required for maintaining both the function and productivity of AM fungi collections.

## Acknowledgements

We thank Ms. Megumi Sakaguchi and Ms. Tamao Nakatani for their technical assistance. The present work was supported by the Cabinet Office, Government of Japan, Cross-ministerial Moonshot Agriculture, Forestry and Fisheries Research and Development Program, “Technologies for Smart Bio-industry and Agriculture” (funding agency: Bio-oriented Technology Research Advancement Institution), JST CREST (JPMJCR16O2), and the JST-Mirai Program (JPMJMI20C7) to YI.

## Conflict of interest

The authors declare that they have no competing interests.

## Notes

### Competing Interest Statement

The authors have declared no competing interest.

